# Integrative analysis of *in vitro* strategies to induce human vascularized cerebral organoids using single-cell RNA sequencing data

**DOI:** 10.1101/2023.02.11.528152

**Authors:** Yuya Sato, Toru Asahi, Kosuke Kataoka

**Author notes:** Correspondence (K.K), (T.A.).

## Abstract

Cerebral organoids are three-dimensional *in vitro* cultured brains that mimic the function and structure of the human brain. One of the major challenges for cerebral organoids is the lack of functional vasculature. Without perfusable vessels, oxygen and nutrient supplies may be insufficient for long-term culture, hindering the investigation of the neurovascular interactions. Recently, several strategies for the vascularization of human cerebral organoids have been reported. However, the generalizable trends and variability among different strategies are unclear due to the lack of a comprehensive characterization and comparison of these vascularization strategies. In this study, we aimed to explore the effect of different vascularization strategies on the nervous system and vasculature in human cerebral organoids. We integrated single-cell RNA sequencing data of multiple vascularized and vascular organoids and fetal brains from publicly available datasets and assessed the protocol-dependent and culture-day-dependent effects on the cell composition and transcriptomic profiles in neuronal and vascular cells. We revealed the similarities and uniqueness of multiple vascularization strategies and demonstrated the transcriptomic effects of vascular induction on neuronal and mesodermal-like cell populations. Moreover, our data suggested that the interaction between neurons and mesodermal-like cell populations is important for the cerebrovascular-specific profile of endothelial-like cells. This study highlights the current challenges to vascularization strategies in human cerebral organoids and offers a benchmark for the future fabrication of vascularized organoids.

## Introduction

Pluripotent stem cell-derived human cerebral organoids are self-organized, three-dimensional *in vitro* cell cultures that mimic the neurodevelopmental processes, organization, and neural activity of the human cerebral cortex. Advancing methods for generating cerebral organoids have provided unprecedented opportunities for understanding neural development, evolution, and disease [1]. Moreover, in the recent coronavirus disease 2019 pandemic, human organoid models have shown promising outcomes in understanding the pathogenesis of the disease, which potentially served as a key to the development of therapeutic agents against severe acute respiratory syndrome coronavirus 2 infections [2,3]. Despite the rapid growth in this field, the development of cerebral organoids is still in its infancy, with several limitations hindering its broader applications and greater impact. One of the major challenges is that cerebral organoids lack vascular systems [4,5]. In the absence of a perfusable vascular network, cerebral organoids rely solely on passive diffusion to exchange nutrients, oxygen, and toxic metabolites. The lack of vascular systems also limits the size of organoids and triggers apoptotic and/or necrotic cell death in the core of organoids [1]. Furthermore, the differentiation of neural progenitor cells is prevented in organoids lacking vasculature [6].

The cerebral vasculature is an uninterrupted arbor-like network of blood vessels comprising diverse cell types, including endothelial cells (ECs), pericytes, and vascular smooth muscle cells [7]. These cells coordinate to support brain homeostasis in a variety of ways: providing oxygen, energy metabolites, and other nutrients to the brain; removing by-products of brain metabolism; preventing the entry of circulating toxins; and modulating immune responses [8,9]. During the development of the nervous system, the vascular system contributes to the proper formation and function of the central nervous system and vice versa [10]. Furthermore, the vascular system controls the proliferation and differentiation of neural progenitor cells through the supply of oxygen and nutrients and also serves as a scaffold for the migration of neuroblasts and newborn neurons [11,12]. Recent studies have proposed that oligodendrocyte precursor cells (OPCs) require blood vessels as scaffolds for migration and that the interaction between OPCs and ECs supports OPC maturation [13].

Multiple strategies have been proposed for the development of vascular systems in human cerebral organoids. In 2018, two groups independently developed robust vascularization of organoid grafts by *in vivo* transplantation of cerebral organoids into mouse brains. Mansour et al. reported the first vascularization strategy of grafting and maintaining cerebral organoids into immune-deficient adult mouse brains for a long term (180 days), in which the organoids exhibited neuronal differentiation and maturation, gliogenesis, microglial integration, and axonal growth into multiple regions of the host brain [14]. The other group (Daviaud et al.) transplanted cerebral organoids into young mice (postnatal day 8–10) without immunosuppressive agents and observed extensive angiogenesis from the host brain into organoid grafts [15].

Furthermore, several protocols have been developed to generate *in vitro* functional vasculatures in human cerebral organoids, which can be broadly classified into three categories (Fig 1A): (1) co-differentiation with mesodermal progenitors, (2) co-culture with ECs, and (3) assembly of distinct organoids/spheroids. The co-differentiation protocols involve the development of embryonic vasculature, starting with the differentiation of mesoderm-derived angioblasts. This strategy closely mimics the development of nascent vasculature during organogenesis [16]. In this strategy, mesoderm development can be induced along with the ectoderm to produce vascularized cerebral organoids using a proper combination of growth factors or gene engineering. Ham et al. guided vasculogenesis by supplementing organoids with vascular endothelial growth factor and Wnt7a [17]. On the contrary, Cakir et al. induced vascular systems by overexpressing human ETS variant transcription factor 2 (*hETV2*) in some populations of stem cells. In this protocol, pericyte-like cells were observed in vascularized organoids in addition to ECs [16]. ETV2 has been extensively studied and utilized to specify cells to endothelial and hematopoietic lineages after the discovery of its capability to convert human skin fibroblasts into functional ECs [18–20].

**Fig 1.**
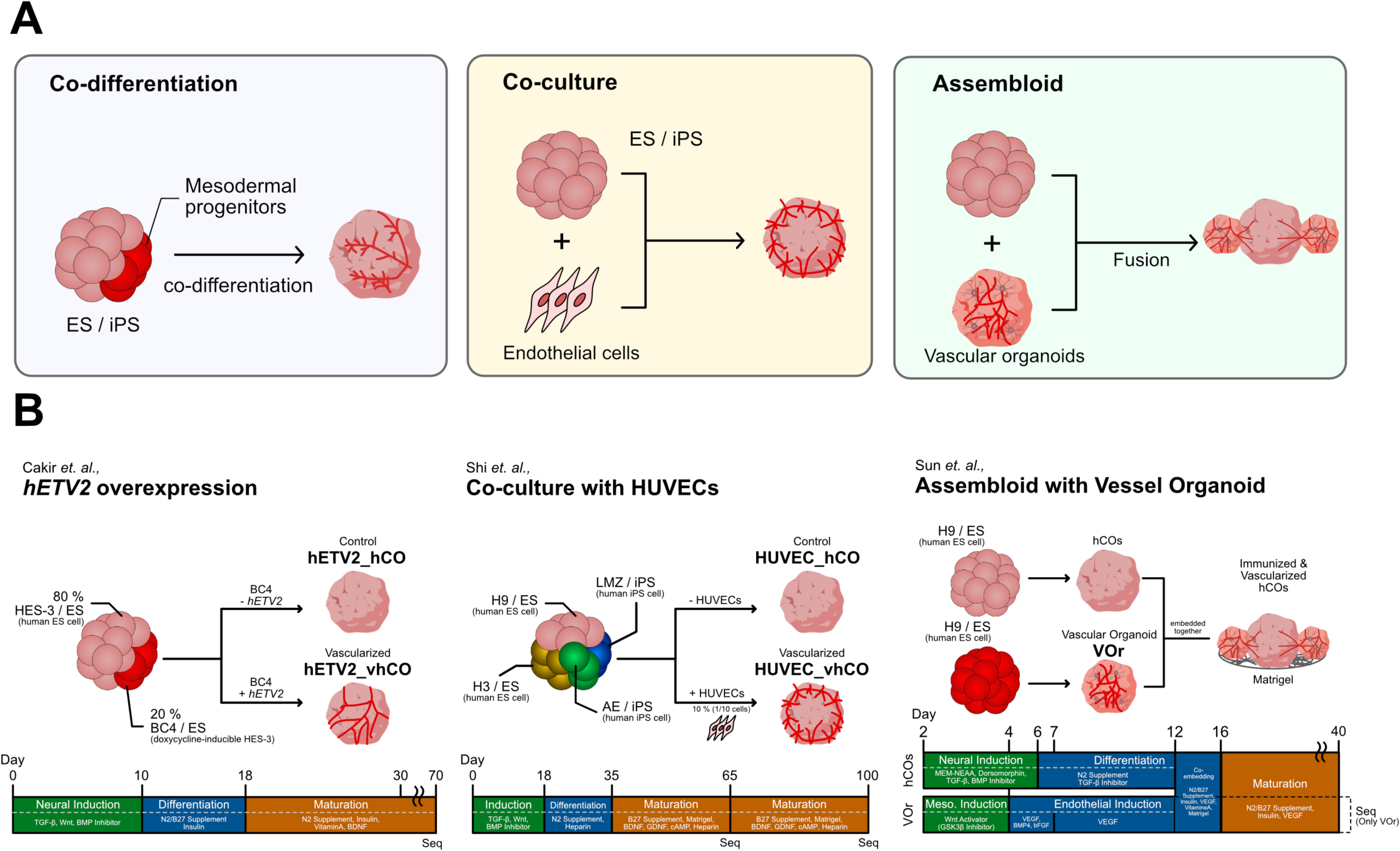
Comparison of vascularization strategies. (A) Three vascularization strategies. Left panel: co-differentiation (differentiation with mesodermal cells), middle panel: co-culture (co-culture with endothelial cells), right panel: assembloid (fusion with vascular organoids). (B) Detailed protocols of the vascularization methods we focused on in this study.

In co-culture protocols, vascularization is achieved by mixing ECs that have already differentiated during organoid formation. There are two main sources of ECs: human umbilical vein endothelial cells (HUVECs) [21] and induced pluripotent stem cell (iPSCs)-derived ECs [21,22]). HUVECs, which originate from the human umbilical vein endothelium, are commonly utilized to vascularize organoids, partly because of the ease with which they are obtained and cultured and were exemplified for the first time by the vascularization in liver buds [23]. Shi et al. used HUVECs to obtain vascularized organoids (termed “vOrganoid” in their study) and successfully cultured them for more than 200 days [21].

The assembly of distinct organoids/spheroids, or assembloids, involves docking cerebral organoids with vascular organoids/spheroids, which are separately differentiated from human embryonic stem cells or iPSCs [24,25]. Unlike other vascularized cerebral organoids, the assembloid proposed by Sun et al. achieved simultaneous immunization and vascularization [24]. Because blood vessels exhibit large heterogeneity of functions among tissues [26], Sun et al. induced specific cerebrovascular features in their vascular organoids by adding neurotrophic factors during the maturation step of vascular organoids, followed by the fusion of brain and vascular organoids [24].

The studies described above proposed different strategies for inducing vascularization in human cerebral organoids, in which the functions, structures, cellular compositions, and cell-cell interactions in the organoids have been independently analyzed. However, no studies have synthetically compared the impacts of these vascularization methods on the diverse cells that comprise vascularized cerebral organoids. Here, we evaluated multiple vascularized human cerebral organoids generated by different strategies through an integrated comparison of single-cell RNA sequencing (scRNA-seq) data available in public datasets and fetal brains. The present study provides insights into the effects of multiple vascularization strategies on cell type differentiation and transcriptomic profiles in neuronal and vascular cells. The findings of this study will provide a benchmark for the fabrication of vascularized cerebral organoids in the future.

## Results and discussion

### Dataset description

We collected publicly available scRNA-seq datasets of human cerebral organoids, which comprised the following three samples: (1) *hETV2* knock-in vascularized organoids [16], (2) HUVEC-co-cultured organoids [21], and (3) vascular organoids (VOr) used for the assembloid [24] (Table 1, S1A Fig). The first two samples ((1) and (2)) included nonvascularized organoids as controls [16,21]. As described in the Introduction section, the vascularization strategies reported in these three studies [16,21,24] were based on three major categories of engineering organoids with functional vasculature (*i.e.*, co-differentiation with mesodermal progenitors, co-culture with ECs, and assembly of distinct organoids/spheroids, respectively) (Fig 1A). The scRNA-seq data from the studies by Cakir et al. and Shi et al. were derived from cerebral organoids that contained induced vasculatures, whereas the data from Sun et al. were derived only from vascular organoids with induced cerebrovascular features for further fusion with cerebral organoids. An overview of the protocols by which these organoids were produced is shown in Fig 1B. We also included scRNA-seq datasets from the human fetal cerebral cortex on 16, 20, 21, and 24 post-conceptual weeks (PCW) [27].

**Table 1.**
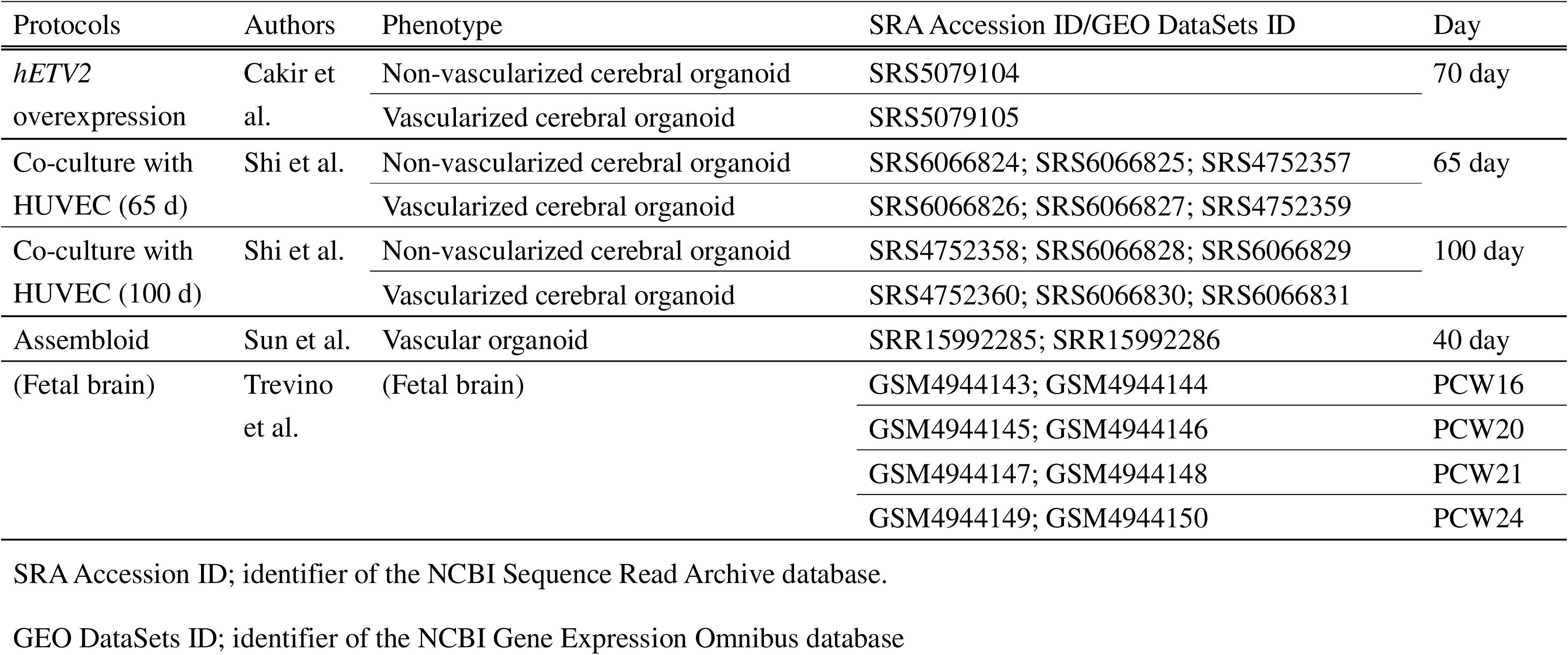
Summary of datasets used in this study.

For pre-processing, we first filtered out low-quality cells from the scRNA-seq datasets (S2A Fig). We then independently analyzed each sample by projecting the cells in each sample into an adjusted 2D space using Uniform Manifold Approximation and Projection (UMAP) and then labeling with any of the 12 cell types based on cell-type-specific marker gene expression, including *SOX2*^+^ progenitor cell (PROG), *TOP2A*^+^/*BIRC5*^+^ proliferating cell (PC), *EOMES*^+^ intermediate progenitor cell (IPC), *TUBB3*^+^/*RBFOX3*^+^ unspecified neuron (Neuron), *GAD1*^+^/*GAD2*^+^ inhibitory neuron (IN), *SLC17A6*^+^/*SLC17A7*^+^ excitatory neuron (EX), *GFAP*^+^/*AQP4*^+^ astrocyte (AS), *OLIG1*^+^/*OLIG2*^+^ oligodendrocyte progenitor cells (OPC), *RBFOX3*^-^/*COL1A1*^+^, *ACTA2*^+^, *RGS5*^+^, *CLDN5*^+^ mesodermal-like cell (MLC), CD53^+^/*CX3CR1*^+^ microglia cell (MGC), *DDIT3*^+^ unfolded protein response cells (UPRC), and unknown cells (Unknown) (Figs 2A, 2B and S4A–S4H). A total of 147,978 cells from multiple samples were integrated based on the feature genes and then visualized using UMAP (Figs 2A and S2D–S2G).

**Fig 2.**
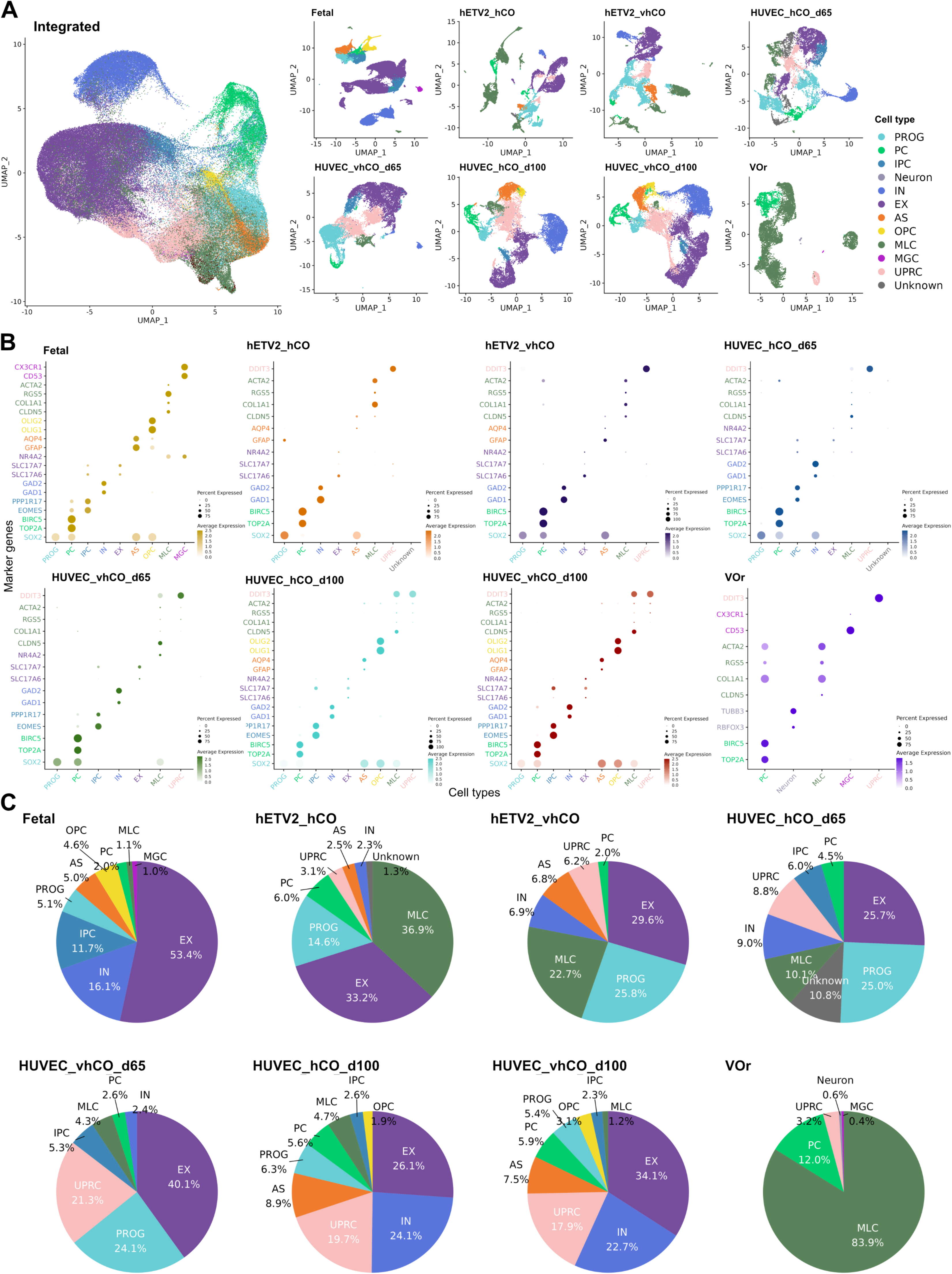
Mapping of scRNA-seq data from non-vascularized and vascularized organoids and fetal brain to UMAP with cell types. (A) UMAP visualization of transcriptomes with cell types. Left panel: UMAP integrated scRNA-seq data for all samples, right panel: UMAP for each sample. Assigned cell types are as follows: PROG; Progenitors (*SOX2*^+^), PC; Proliferating Cells (*TOP2A*^+^, *BIRC5*^+^), IPC; Intermediate Progenitor Cells (*EOMES*^+^, *PPP1R17*^+^), Neuron (*RBFOX3*^+^, *TUBB3*^+^), IN; Inhibitory Neurons (*GAD1*^+^, *GAD2*^+^), EX; Excitatory Neurons (*SLC17A6*^+^, *SLC17A7*^+^), AS; Astrocytes (*GFAP*^+^, *AQP4*^+^), OPC; Oligodendrocyte Progenitor Cells (*OLIG1*^+^, *OLIG2*^+^), MLC; Mesodermal-Like Cells (*CLDN5*^+^, *COL1A1*^+^, *RGS5*^+^, *ACTA2*^+^), MGC; Microglial Cells (CD53^+^, *CX3CR1*^+^), and UPRC; Unfolded Protein Response Cells (*DDIT3*^+^). (B) Expression of marker genes of each cell type in each sample. The cell type is displayed on the x-axis, and the marker gene for each cell type is displayed on the y-axis. The color of the marker corresponds to the color of each cell type. hCO; human cerebral organoids, and vhCO; vascularized hCO. (C) Ratio of cells present in each sample. The ratio is calculated as the percentage of each cell count to the total cells in each sample.

### Cell type compositions in vascularized, non-vascularized, and vascular organoids

The annotation strategy of the 12 cell types is shown in S1B Fig. Each assigned cell type specifically expressed the marker genes (Fig 2B). Moreover, the top 100 genes enriched in each cell type scarcely overlapped, suggesting that the assigned cell types were independent populations, representing separate transcriptomic profiles (S3A Fig). Additionally, genes specifically expressed in PROGs, neuronal cell types (IPC, Neuron, IN, and EX), and MLCs were shared with those expressed in the fetal brain, suggesting that the characteristics of each cell type in the organoids were similar to those of the fetal brain at the transcriptomic level (S3B Fig).

Next, we sought to identify differences in cell composition with vascularization by calculating the percentage of cell type occupancy in each sample. Except for that in VOr, a relatively large occupancy of the following cell types was observed PROG, PC, IN, and EX in all organoids (Fig 2B and 2C). Furthermore, we found that major changes in cell composition were induced by the protocols or days of culture rather than by vascularization (Figs 2C and S2C). For the protocol-dependent cell-composition difference, the organoids in Cakir et al.’s study were characterized by a high ratio of MLC and the presence of AS [16]. In contrast, those in Shi et al.’s study at day 65 of culture were characterized by a low ratio of MLC and the absence of AS. For the day of culture-dependent differences, a prolonged culture period induced AS and OPC in the organoids [21], which might be due to the increased diversity of cell types with the time of growth [28] (Fig 2B and 2C).

Cell populations with non-microglial mesoderm-like expression were assigned to MLCs, which cannot be classified into any cell type in ectodermal lineages such as neurons and astrocytes. Most of the cells in the VOr were classified as MLC with the expression of *CLDN5*, *COL1A1*, *RGS5,* and *ACTA2* (Figs 2B, 2C, and S4H), as VOr solely comprised the vascular system. We also identified MLCs in organoids from Cakir and Sun’s studies (Fig 2B and 2C). The lack of dual-SMAD inhibition (inhibition of BMP and TGF-β) during the generation of cerebral organoids can lead to the formation of mesoderm-derived progenitors [29,30]. Nevertheless, Cakir and Sun’s studies applied LDN-193189 and SB-431542 to inhibit BMP/TGF-β signaling in neural induction processes, suggesting that cerebral organoids innately develop MLCs irrespective of dual-SMAD inhibition or vasculature induction.

Another challenge for human cerebral organoids is the absence of microglia and vascularization [31]. Microglial clusters (MGC) were absent from cerebral organoids in Cakir et al.’s [16] and Shi et al.’s [21] studies, whereas they were found in VOr in the study by Sun et al. [24] (Fig 2C). These results suggest that immunization is also achievable by the vascularization of cerebral organoids, according to the protocol of Sun’s study. In addition, VOr contained a small number of *RBFOX3^+^*/*TUBB3^+^* neuronal cell clusters (Fig 2B and 2C). This cluster lacked the typical EX and IN marker genes. These neurons that developed in the VOr may be, therefore, off-target immature neurons that have differentiated by the supplement 0of N2/B27 from undifferentiated cells during the maturation process.

Overall, we identified the diversity of cell types in vascularized organoids, including newly discovered mesodermal cells, and protocol- or culture-time-dependent heterogeneity of cell types.

### Diverse vascularization protocols trigger different transcriptomic alterations in human cerebral organoids

We next examined whether different vascularization strategies influence the fidelity of organoids to the human fetal brain. To this end, we calculated the correlations of “enrichment score” in each cell type in organoids and the human fetal brain (see Methods). All vascularization protocols improved the correlations in most cell types, suggesting that vascularization generally advances organoid fidelity to the human fetal brain (Fig 3A).

**Fig. 3.**
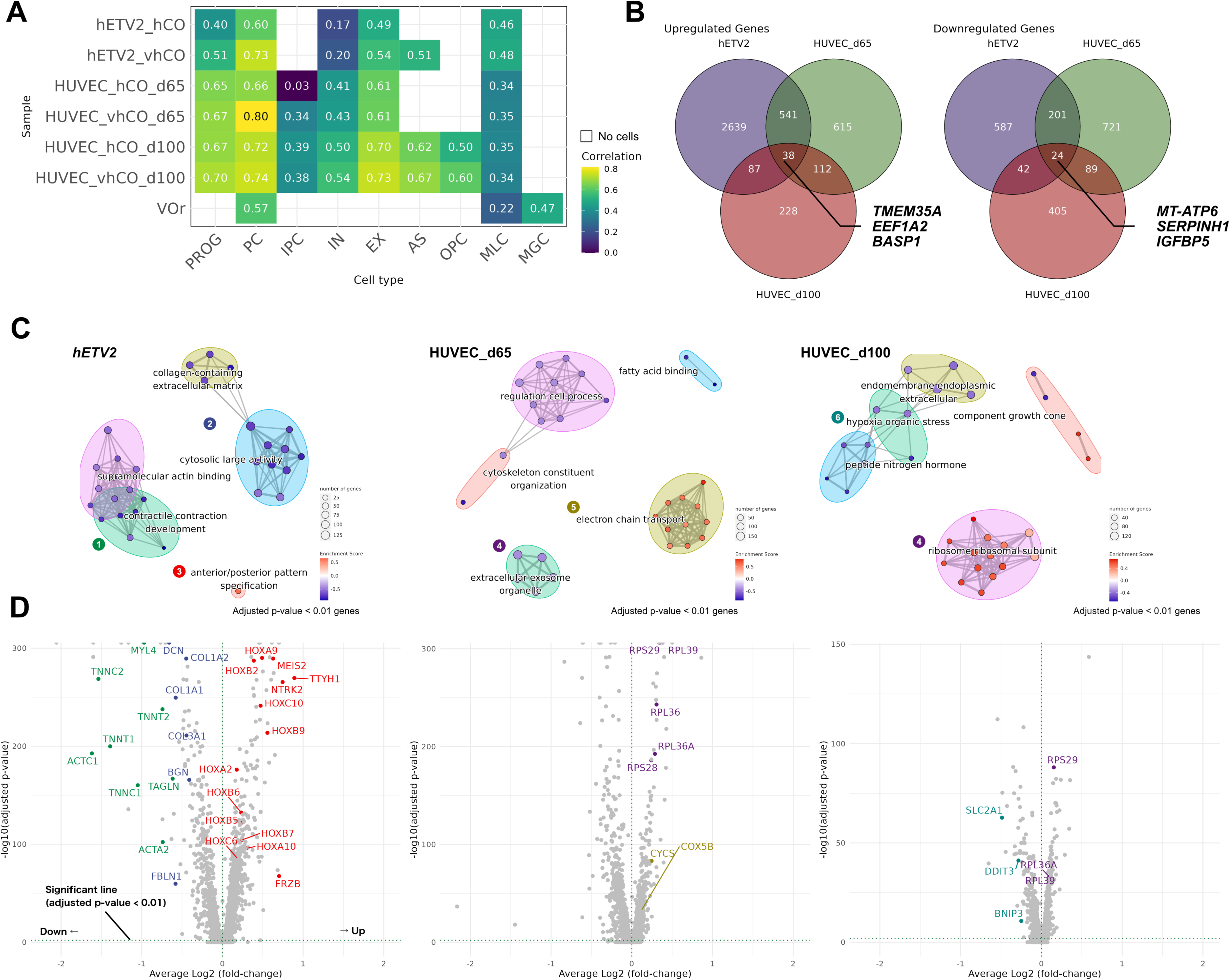
Transcriptomic fidelities in the fetal brain and bulk alteration transcriptomic by vascularization. (A) Similarity of transcriptome profiles in the fetal brain. Higher correlation values indicate higher similarity. Blank cells indicate the absence of cells [Refer to Methods section for the calculation of the correlation values]. (B) Venn diagrams of genes altered by vascularization. Numbers indicate the number of genes changed. Left panel: Venn diagram of upregulated genes, right panel: Venn diagram of downregulated genes. Genes pointing to the center represent examples of genes commonly altered by vascularization in all vascularization protocols. (C) Significantly altered gene ontology (GO) terms (p-value < 0.05) predicted by gene set enrichment analysis (GSEA) of genes significantly altered by vascularization (corrected p-value < 0.01). The color of the nodes represents the enrichment score (the amount of change calculated by GSEA). (D) Volcano plot of genes with altered expression induced by vascularization. The average log_2_ fold-change (FC) is shown on the x-axis and the −log_10_ (corrected p-value) is on the y-axis. The color of the gene names corresponds to the color of the GO terms in (C).

Nevertheless, the transcriptomic similarities of MLCs in the VOr were not as high (0.22; Fig 3A) as those of the organoids in previous studies [16,21], suggesting that treatment with neurotrophic factors to induce cerebrovascular features in the VOr was insufficient to reproduce brain-specific MLC.

Next, we investigated the bulk transcriptomic alterations triggered by different vascularization methods. The calculation of the number of genes whose expression levels were significantly altered by vascularization identified 4260 upregulated and 2449 downregulated genes. Overexpression of *hETV2* upregulated 3305 genes, whereas co-culture with HUVECs upregulated only 1306 genes at 60 days and 465 genes at 100 days (Fig 3B). These differences suggested that *hETV2* activates diverse biological processes. The number of genes commonly upregulated and downregulated in the three samples was 38 and 24, respectively (Fig 3B and STable 1). The commonly upregulated genes were related to neuronal development and axonogenesis, including *TMEM35A*, *EEF1A2*, and *BASP1* [32–34]. The commonly downregulated genes were related to neurotoxicity, such as *MT-ATP6*, *SERPINH1*, and *IGFBP5* [35–37]. These results imply that vascularization increases neural activity and suppresses cellular stress, although a few processes are consistently altered across protocols.

We predicted overrepresented biological processes using gene set enrichment analysis (GSEA) based on the significant differentially expressed genes (adjusted *p*-value < 0.01) between nonvascularized and vascularized organoids (Figs 3C, 3D, S4I and STable2). GSEA revealed that overexpression of *hETV2* downregulated collagen-containing extracellular matrix, actin binding, and contractile contraction development, while it upregulated anterior/posterior pattern specification (Fig 3D). Downregulated genes included extracellular matrix-related genes, such as collagen family genes (*COL1A1*, *COL1A2, COL3A1*), fibulin (*FBLN1*), and proteoglycans (*BGN, DCN*); actin binding-related genes, including *ACTA2* and *ACTC1* (actin family genes); and muscle function-related genes, including troponin family genes (*TNNT1*, *TNNT2*, *TNNC1*, *TNNC2*), myosin family genes (*MYLIP* and *MYL4*), and transgelin (*TAGLN*). ETV2 is known to directly reprogram fibroblasts or muscle cells into vascular ECs [18,38]. The observed downregulation of the extracellular matrix and muscle development by *hETV2* overexpression might be due to its trans-differentiation capability. GSEA also revealed that the anterior/posterior pattern specification process was upregulated, in which the expression of homeobox family genes (*HOXA2, HOXB2, HOXB5, HOXB6, HOXB7, HOXC6, HOXA9, HOXA10, HOXC10*) was enhanced as previously reported [16]. In contrast, the genes encoding the electron transport chain (*COX5B*, *CYCS*) and ribosomal proteins (*RPL39, RPS29, RPL36, RPL37A, RPS27, RPL36, RPL36A, RPS28, RPL38, RPS27L, RPS24*) were upregulated in HUVEC co-cultured with organoids (Fig 3A and 3B), which indicated the availability of sufficient nutrient supply by HUVEC vascularization. Moreover, after 100 days of HUVEC culture, the expression of hypoxia markers *BNIP3* and *SLC2A1* (protein: GLUT-1) and an unfolded protein response marker *DDIT3* was significantly downregulated, indicating that HUVEC co-culture suppressed the hypoxic stress. These results suggest that vascularization with HUVECs potentially improves oxygen and nutrient supply, leading to cellular stress suppression, whereas *hETV2* overexpression regulates cell differentiation capacity and regional patterning. Taken together, these studies imply that vascularization increases the fidelity of cell differentiation in cerebral organoids strategy-dependently.

### Different vascularization protocols uniquely influence the transcriptome of neuronal populations in cerebral organoids

The cerebral organoids in the Cakir’s (*hETV2* overexpression) and Shi’s (HUVEC co-culture) studies produced three neuronal subtypes: EX, IN, and intermediate progenitor cells (IPC).

Therefore, we set out to characterize the effect of these vascularization methods on the transcriptomic profile, with a focus on these neuronal populations. Datasets derived from VOr were excluded from this analysis because they contained few neuronal cells. First, we extracted and re-clustered neural cells and then projected them into UMAP (S5A Fig). The analysis revealed *hETV2* overexpression-mediated vascularization increased the proportion of IN (*GAD1*^+^/*GAD2*^+^) from 6.6% to 18.9% (Fig 4A). In contrast, vascularization achieved by co-culture with HUVECs increased the proportion of EX (*SLC17A7*^+^/*KCNJ1*^+^) from 66.1% to 83.9% on 65 days and from 49.5% to 57.7% on 100 days (Fig 4A). These results indicated vascularization methods also alter the proportions of neuronal subtypes.

**Fig. 4.**
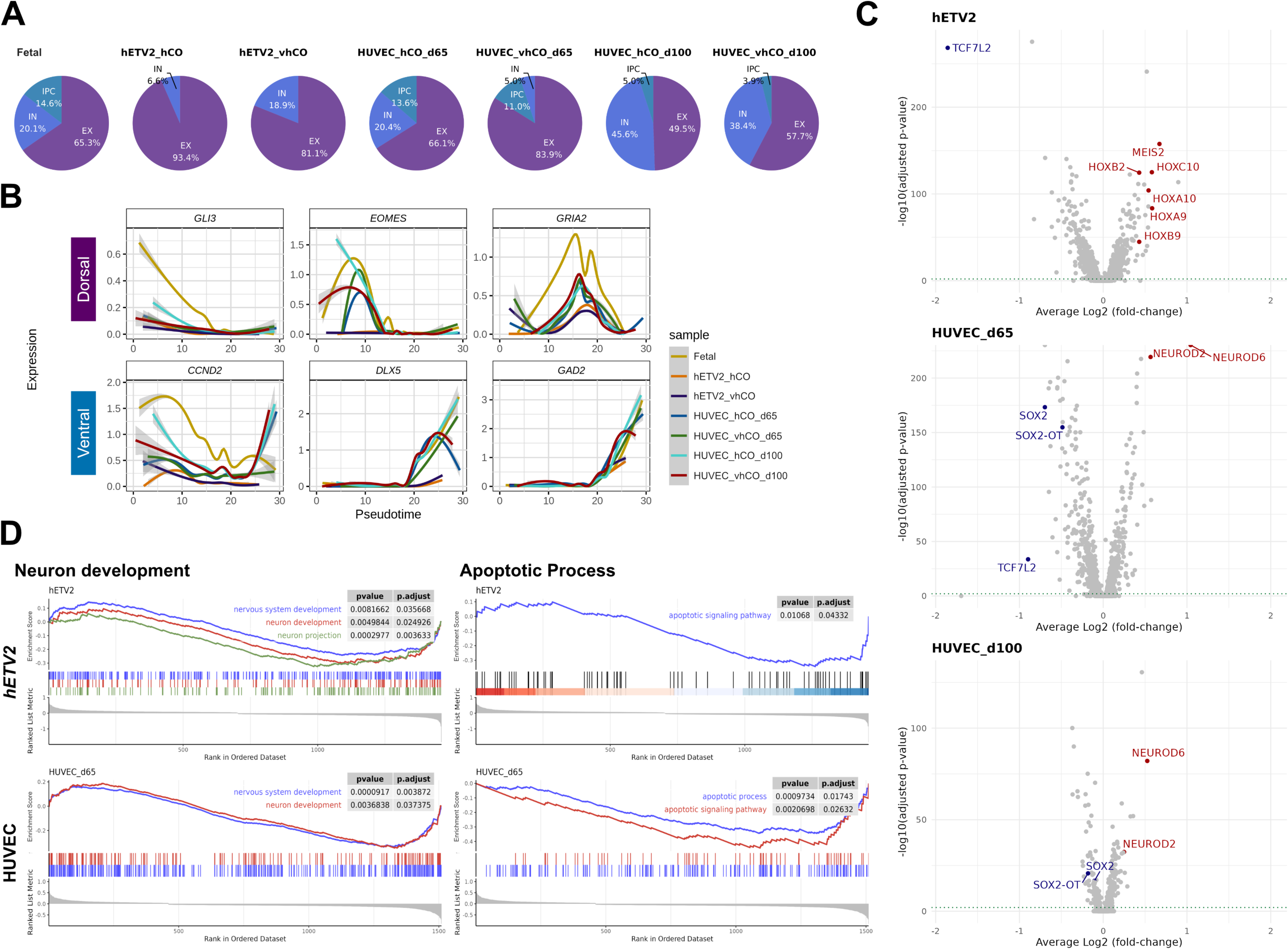
Protocol-dependent differences in transcriptomic profiles in neuronal cell populations. (A) Percentage of neuronal subtypes in each sample. (B) Changes in expression levels of genes characteristic of dorsoventral development along pseudotime. The pseudotime was calculated with predicted differentiation trajectories of the following cells: PROG, PC, IPC, Neuron, IN, EX, AS, and OPC. (C) Volcano plot of gene expression altered by vascularization in neurons. (A) (D) GSEA plots of GO terms predicted from the significantly altered (adjusted p-value < 0.01) gene groups. Left panel: GO terms involved in neurodevelopment (nervous system development, neuron development, neuron projection), right panel: GO terms involved in apoptosis (apoptotic signaling pathway). In HUVECs co-culture samples for 100 days, GO terms were not predicted because of the low number of genes altered by vascularization.

In the developing fetal brain, cortical EXs are generated locally in the dorsal forebrain, whereas INs are generated in the ventral forebrain and migrate to the cortex [39]. To uncover the underlying molecular mechanisms of the differences in neuronal subtype differentiation, we analyzed the changes in the expression of dorsal- and ventral-specific neurogenic genes over time. To predict the pseudotime, ectodermal cells (IPC, Neuron, EX, IN, AS, and OPC) plus PROG and PC were extracted and re-clustered, followed by the construction of pseudo-differentiation trajectories. Vascularization had insignificant effects on the temporal expression patterns of most neurogenic genes, except for *EOMES* (Fig 4B). *EOMES* (protein: Tbr2) is specifically expressed in IPCs, which are derived from radial glial progenitors and neural stem cells of the developing cortex, and serve as excitatory neurogenic progenitors [40,41]. As shown in Fig 4B, *EOMES* expression was enhanced in a specific developmental time window in the 65-day HUVEC co-cultured organoid [21], whereas *hETV2*-overexpressed organoids [16] lacked *EOMES*. Because IPC is believed to contribute to cortical expansion in primates [42,43], the upregulation of *EOMES* at 65 days possibly induced organoid expansion. Concordantly, Shi et al. demonstrated a rapid expansion in the size of organoids induced by vascularization around 65–70 days [21]. These findings support the importance of vascularization in neurogenesis in human cerebral organoids. However, it cannot be ruled out that these results may arise from differences in culture periods rather than the differences in vascularization protocols.

Next, to unbiasedly identify transcriptomic changes associated with vascularization, we comprehensively analyzed genes differentially expressed by vascularization in neuronal populations (Fig 4C and STable3). The upregulated genes in *hETV2*-overexpressed organoid contained many homeodomain-containing transcription factors, such as *HOXA9*, *HOXA10*, *HOXB2*, *HOXB9*, *HOXC10,* and *MEIS2*, as observed in the previous analysis of all cell types. In organoids co-cultured with HUVEC, neuroid family genes such as *NEUROD2* and *NEUROD6*, which are proneural basic helix-loop-Helix (bHLH) transcription factors responsible for neuronal differentiation and specification [44], were significantly upregulated (Fig 4C). These gene alterations were consistent with the upregulation of GO terms involved in “Neuron development” based on GSEA (Fig 4D, left panel). Consistent with our finding that HUVEC co-culture enhanced EX proportions (Fig 4A), *NEUROD2/6* is important for glutamatergic function in cortical neurons [45]. In addition, in organoid co-culture with HUVEC protocol*, SOX2* and long noncoding RNA *SOX2-OT* were downregulated in 65-day culture, whereas only *SOX2-OT* was downregulated in 100-day culture protocols (Fig 4C).

Furthermore, *TCF7L2* was suppressed in *hETV2*-overexpressed and HUVEC co-cultured (65-day) organoids (Fig 4C). Decreased expression of these genes has been linked to reduced Wnt activity and suppression of the proliferation of radial glial cells and intermediate progenitors [46]. Consistently, the proportion of *EOMES*^+^ IPCs was reduced by HUVEC co-culture from 13.6% to 11.0% in the 65-day culture and from 5.0% to 3.9% in the 100-day culture (Fig 4A). GSEA analysis predicted that alterations in neurodevelopmental processes as well as downregulation of biological processes, are involved in apoptosis in organoid neurons, irrespective of the vascularization protocols. This indicates that vascularization generally prevents neuronal cell death. Altogether, these findings suggest that vascularization regulates the balance between neuronal cell differentiation and progenitor cell proliferation in cerebral organoids. However, understanding the underlying molecular mechanisms require further study.

### Characterization of vascular-like cells that develop in human cerebral organoids

Finally, we focused on the MLC population, which are neither neural (*RBFOX3*^+^), proliferative (*TOP2A*^+^), nor glial cells (*GFAP^+^, AQP*^+^, and *OLIG2*^+^), expressing genes characteristic of mesodermal-derived cells (Fig 1B). To further investigate these cells, we isolated them from the combined dataset (Fig 5A). These cells expressed genes characteristic of cells that comprise blood vessels, such as *CLDN5*, *ACTA2*, *COL1A1,* and *RGS5* (Fig 5B). The isolated cell population was re-clustered at a resolution of 0.3, yielding 11 clusters (S6A Fig). The resolution was determined using a “clustree” package in Seurat, illustrating the cluster relationships at multiple resolutions (0 to 1 in 0.1 increments) (S6B Fig). Cell types were assigned to these clusters according to GSEA based on the unbiasedly determined marker gene expression (Figs 5C and S6C, S6D). These cell types include vascular-like cells, such as ECs, fibroblasts, and mural cells (pericytes and smooth muscle cells).

**Fig. 5.**
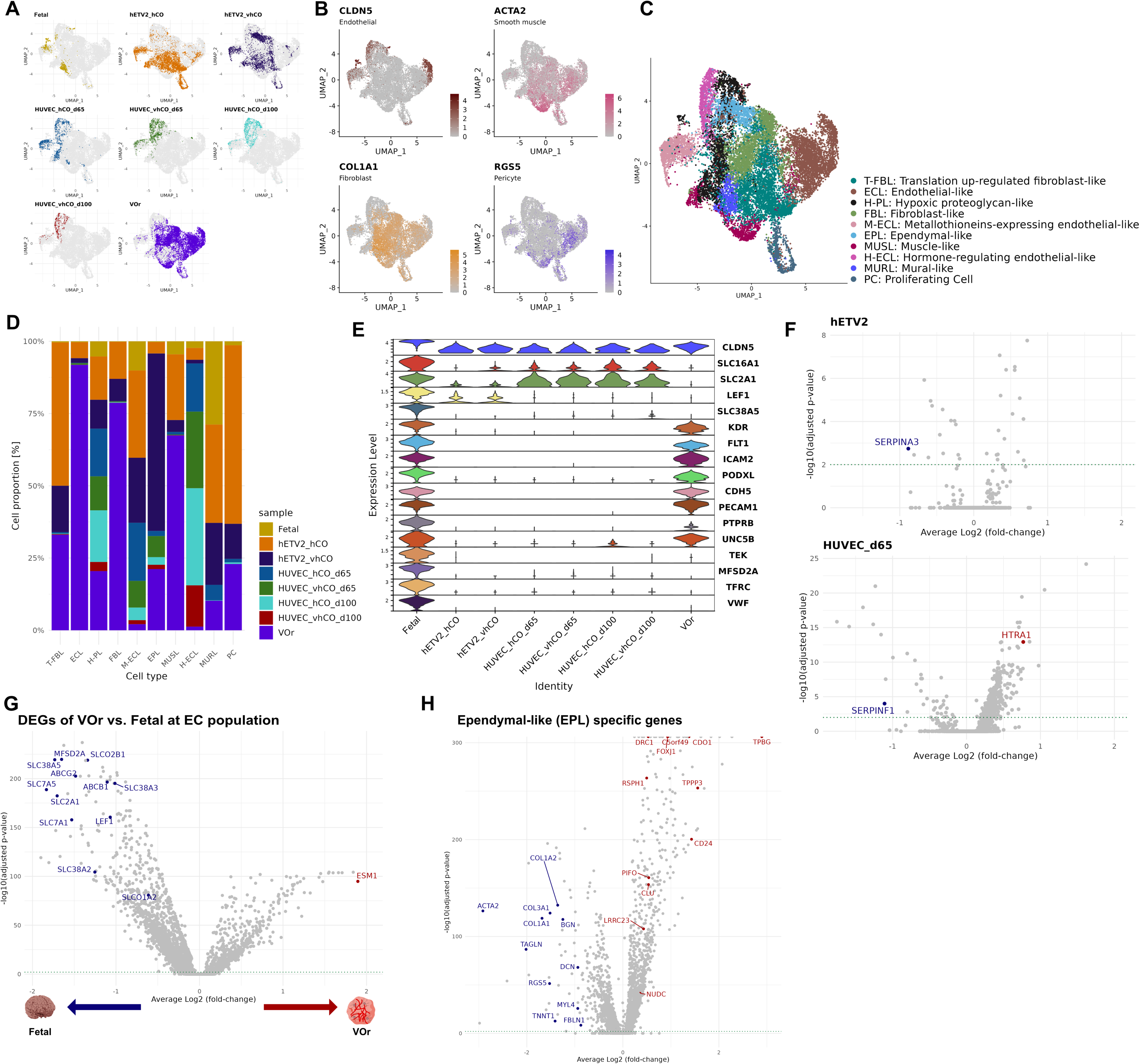
Characterization of transcriptomic profile alterations with vascularization in mesodermal-like cell (MLC) subtypes. (A) UMAP for each sample of MLC. (B) Marker expression of vascular cells (*CLDN5*, endothelial cells*; ACTA2*, smooth muscle cells; *COL1A1*, fibroblast; *RGS5*, pericytes). (C) UMAP assigned to cell types based on unbiased gene expression profiles [See S6A Fig for clusters to which cell types are assigned; S6C and S6D Fig show the annotation]. (D) Proportion of cells in each sample per cluster. (E) Endothelial cell marker expression. (F) Volcano plot of genes altered by vascularization in the *CLDN5*^+^ endothelial-like (ECL) cells. The data for organoids co-cultured with HUVECs for 100 days are not shown due to the low number of genes altered by vascularization. (G) Differentially expressed genes in VOr ECL cells (*CLDN5*^+^, *KDR*^+^, *FLT1*^+^) compared to endothelial cells in the fetal brain. The larger the averaged log_2_(FC), the more enriched in VOr, and vice versa. (H) Gene expression specific to an ependymal-like (EPL) cluster. Genes colored in red represent highly expressed genes in EPL compared to other clusters, and genes colored in blue represent less expressed genes.

The three endothelial-like (ECL) cell types, ECL, metallothionein-expressing ECL (M-ECL), hormone-regulating ECL (H-ECL), commonly expressed *CLDN5,* a component of brain endothelial-specific tight junctions known as a robust EC marker (Fig 5B and 5E) [9,47]. M-ECL and H-ECL expressed unique transcriptomic profiles, such as metallothioneins and thyrotropin-releasing hormone (TRH), respectively, in addition to *CLDN5*. Metallothionein contributes to the detoxification of excess heavy metals and the removal of reactive oxygen species, while TRH regulates hormone secretion in the brain [48,49]. M-ECL and H-ECL were present predominantly in cerebral organoids (*hETV2*-overexpressed and HUVEC co-cultured organoids) and, to a lesser extent, in the fetal brain and VOr (Fig 5D). In contrast, VOr contained abundant ECL with the expression of *PECAM1* (protein: CD31) and *FLT1* (protein: VEGFR1), which are important for defining endothelial properties [9,50,51]. To further highlight the heterogeneity in ECL cells across different protocols, we characterized the expression of commonly referenced marker genes characteristic of cerebral vascular ECs. To achieve this, we extracted *CLDN5*^+^ ECL cells from all MLCs and profiled their marker genes (Fig 5E). Genes encoding solute carrier proteins in human brain capillary ECs, such as *SLC16A1* and *SLC2A1*, are barely expressed in VOr [52]. Furthermore, *LEF1*, a transcription factor controlling the blood-brain barrier (BBB)-specific gene expression repertoire, is expressed only in the fetal brain and part of the cerebral organoids [53]. In contrast, *CLDN5^+^* ECL cells in the cerebral organoids lacked the expression of *PECAM1* and VEGFRs (*KDR* and *FLT1*) (Fig 5E). To ascertain whether vascularization in these cerebral organoids improves their ECL cell characteristics, we explored genes whose expression was significantly altered by vascularization in *CLDN5*^+^ ECL cells (Fig 5F and S4 Table), which identified a few genes significantly altered by vascularization (Fig 5F). On the contrary, several cerebrovascular-associated genes were significantly altered by vascularization in the *CLDN5^+^* cell population. For example, the expression of *SERPINF1*, which belongs to the serpin family and has antiangiogenic activity [54], decreased in HUVEC-co-cultured (65-days) organoids. Furthermore, in *the hETV2*-overexpressed organoid-derived *CLDN5^+^* cell population, the expression of *SERPINA3*, which negatively regulates angiogenesis and inflammation, was reduced [55]. In addition, *HTRA1*, a causative gene for cerebral small-vessel disease [56,57], was upregulated in HUVEC co-cultured (65 days) organoids. These findings suggest that ECL cells in cerebral organoids possess properties specific to cerebrovascular cells but lack sufficient properties as vascular ECs. Furthermore, vascularization may induce the expression of several gene profile characteristics of the cerebrovascular system. However, it may not lead to the acquisition of full EC properties for ECL cells. To characterize ECL cells in the VOr, we identified differentially expressed genes between ECL cells in the VOr and fetal brain ECs. The expression of ESM1, a protein synthesized and secreted by ECs in the lungs and kidneys, was elevated in VOr [58] (Fig 5G and S5 Table). Conversely, the expression of BBB-related genes, such as a regulator of BBB permeability (*MSFDA2A*), BBB-specific efflux (*ABCB1* and *ABCG2*), and influx (*SLCO1A2* and *SLCO2B1*) transporters [59,60] was reduced (Fig 5G). These findings suggest that the EC population of VOr expresses the core genes as real ECs but fails to reproduce the characteristics of cerebral blood vessels despite treatment with neurotropic reagents N2 and B2 at the late stage of maturation.

We also found non-vascular cells in the MLCs, such as proliferating cells (PC) and ependymal-like cells. Ependymal cells in the human brain control cerebrospinal flow to efflux waste products [61]. To investigate the nature of the EPL in organoids in detail, we performed a differential expression test between the EPL cluster and the other clusters. The EPL showed enhanced expression of ciliogenesis-related genes (*TPBG*, *CDO1*, *CD240*, *DRC1*, and *C5orf49*) [62] and suppressed expression of perivascular-like cells marker genes (*COL1A1*, *ACTA2,* and *TAGLN*) (Fig 5H and S6 Table). We also found that EPL upregulated *FOXJ1*, *CLU*, *PIFO*, *DYMLRB2*, *LRRC23*, *RSPH1*, *TPPP3*, and *NUDC* (Fig 5H), the marker genes of ependymal cells [63–65]. Moreover, the overexpression of *hETV2* remarkably increased the population of EPL. These results suggest that *hETV2* overexpression induces vasculogenesis as well as ependymal-like cell development.

Collectively, ECs could be present in organoids regardless of vascularization, whereas vascularization induced alterations in functional proteins in these cells. There is a trade-off relationship between the expression levels of genes critical to ECs (*e.g.*, *PECAM1* and *FLT1*) and those expressed in a BBB-specific manner (*e.g.*, *SLC2A1*), in which the balance of expression of both types of genes depends on vascular induction strategies.

## Conclusions

Various cerebral organoid vascularization strategies improve the survivability and reproducibility of cerebral organoids. The increasing volume of publicly available scRNA-seq datasets of human cerebral organoids has enabled comparative studies to identify generalizable trends and/or variability among individual studies. In this study, we systematically characterized the single-cell transcriptome of vascularized cerebral and vascular organoids with induced cerebrovascular features. Our results reveal how vascular induction has transcriptomic effects on neuronal and mesodermal-like cell populations. Moreover, our data suggest that the interaction between neurons and mesodermal-like cell populations is important for the cerebrovascular-specific profile of endothelial-like cells. The benchmarks we constructed suggest that diverse vascularization strategies have issues that need to be resolved. In future studies, detailed analysis of the induced vasculature using techniques such as spatial transcriptomics and vasculature-targeted scRNA-seq will be required to precisely evaluate vascular cell types and their functional aspects.

## Methods

### Data collection

We collected the FASTQ or BAM-formatted files from the NCBI Short Read Archive for the 10x Genomics platform. The downloaded files were converted to FASTQ files using “fastq-dump” (v2.11.0) or “bamtofastq” (v1.3.2). The FASTQ files were mapped to the human reference genome (GRCh38, v1.2.0) using the Cell Ranger count function (v6.0.1) with the default parameters. Multiple sequences in the same experiment were pooled using the “aggr” function of Cell Ranger. For samples without 10x Genomics platform data, we obtained post-mapped cell-gene matrices from the Gene Expression Omnibus (GEO) database.

### Pre-processing

We quality-controlled the cell-gene matrix output from the Cell Ranger analysis pipeline named “filtered_feature_bc_matrix” to exclude low-quality cells. Low-quality cells exhibited 1) high mitochondrial expression, 2) low feature mRNA expression, or 3) multiplet profiles [61]. Therefore, we excluded low-quality cells from the scRNA-seq libraries of cerebral and vascularized cerebral organoids and fetal brain under the following conditions:1) organoid, >5%; fetal brain, >10%; and 2) less than 1000 feature genes. A multiplet is defined as an artificial single-cell profile produced by two or more cells containing the same barcode. To filter the multiplets, we predicted and eliminated multiplets using Single-Cell Remover of Doublets (Scrublet v0.2.3) with default parameters [66].

### Data integration

Pre-processed matrices from multiple samples were integrated and clustered using Seurat (v4.0) in R (v4.1.3) environment. Variations in technical factors, including the sequencing depth in samples obtained from different experiments, lead to batch effects [67]. To eliminate this variation, we used the “SCTransform” function to normalize and variance-stabilize gene counts. Next, feature genes for integration were selected using the “SelectIntegrationFeatures” function and normalized by the “PrepSCTIntegration” function. Then, we identified the integration anchors of all sample data using the “FindIntegrationAnchors” function and integrated all assays with these anchors using the “IntegrateData” function. The integrated gene expression was compressed to 30 dimensions using the “RunPCA” function, and then the feature genes from 1 to 20 dimensions were k-mer clustered using the “FindNeighbors” and “FindClusters” functions. The compression dimensions were selected using the “ElbowPlot” function.

### Cell type determination

Cell types were determined using the following steps: 1) normalization and scaling, 2) dimensionality reduction, 3) clustering, and 4) cell type assignment using marker genes for each sample. These processes were performed differently from the “Data Integration” process to correctly assign cell types in the sample even under varying profiles among samples. The determined cell types were represented one-to-one with cell barcodes for use in all subsequent analyses. Each step was implemented according to Seurat’s Vignette by normalization and scaling with the “SCTransform” function, PCA and UMAP dimensionality reduction from 1 to 20 dimensions, clustering with “FindNeighbors” and “FindClusters,” and cell type determination using the reported marker genes.

### Differentially expressed gene analysis

To capture changes in gene expression profiles, gene sets varying among two conditions (e.g., vascularized organoid and fetal brain) were identified by the Wilcoxon Rank Sum test using the “FindMarker” function in the Seurat package. The “FindMarker” function was implemented with the target condition (ident.1 parameter) and the comparison condition (ident.2 parameter). From the calculated genes, only gene sets satisfying the condition (adjusted p-values with Bonferroni correction) < 0.01 were extracted and used in the subsequent analysis, except for volcano plots. Volcano plots were plotted as average log_2_FC on the x-axis and -log_2_(adjusted p-value) on the y-axis with full gene sets. Among the gene sets of interest, upregulated genes are highlighted in red, and downregulated genes are highlighted in blue. In addition, a threshold line (adjusted p-value < 0.01) is represented by the green dotted line.

### Sample correlation with cell type

To determine the genetic similarity of human cerebral organoids to the fetal brain, we calculated the correlations of unbiased marker genes in each cell type of each organoid and fetal brain. First, we calculated marker genes, which are cell type-specific upregulated genes, with the Wilcoxon Rank Sum test using the “FindMarker” function in the Seurat package (see **Differentially expressed gene analysis**). Then, we calculated Pearson’s correlations for log_2_FC of marker genes under (adjusted *p*-values with Bonferroni correction) < 0.01 to define similarity.

### Gene set enrichment analysis (GSEA)

We performed GSEA to predict biological processes from differentially expressed genes (See **Differentially expressed gene analysis**) using the “clusterProfiler” package (v4.2.2) in the R environment. First, we predicted pathways using a “gseGO” function with default parameters under the condition (adjusted *p*-values with Bonferroni correction) < 0.01. Second, we computed the similarity matrix between GO terms using the “pairwise_termsim” function of the “enrichplot” package (v1.14.2) and visualized gene sets as a network using the “emapplot” function. The color range was determined by the enrichment score, which indicated the predicted variability of each term.

### Pseudotime analysis

To determine the changes in gene expression during the differentiation time course, cell trajectory lineages and differentiation pseudotime were calculated using the slingshot package (v2.2.0). The 2D UMAP coordinate matrix was extracted from the integrated Seurat object to execute a trajectory of slingshot analysis with all Seurat clusters and the root cluster number as the input. The root cell cluster was defined as *TOP2A*^+^ proliferating cells with the highest expression of *SOX2*. A minimum spanning tree on the cluster is constructed by the “getLineages” function to identify the phylogenetic relationships across the cluster. We then inferred the pseudotimes by fitting the principal curves using the “getCurves” function. The calculated pseudotimes were averaged using the “averagePseudotime” function of the “TrajectoryUtils” package to calculate the pseudotime at each lineage.

### Gene expression along the pseudotime course

To characterize differences in gene expression during differentiation progression, we plotted gene expression along the pseudotime on a differentiation trajectory. The pseudotime was calculated by trajectory analysis using a slingshot and then averaged for each differentiation trajectory. Gene expression along the pseudotime was fitted by a loess approximation using the “geom_stat” function in the ggplot2 package (v 3.3.6).

## Supporting information

STable 1

STable 2

STable 3

STable 4

STable 5

STable 6

## Acknowledgments

This study was supported by grants from JSPS KAKENHI (19K20196 and 22K17815 to KK).

## Author contributions

Conceptualization: Y.S. and K. K.; data curation: Y. S.; formal analysis: Y.S.; funding acquisition: K.K. and T. A.; investigation: Y.S. and K. K.; methodology: Y. S.; project administration: K.K.; software: Y.S.; supervision: K. K. and T. A.; validation: K. K.; visualization: Y. S.; writing – original draft: Y. S. and K. K.; writing – review and editing: Y. S. and K. K. All authors have read and agreed to the published version of the manuscript.

## Abbreviations

BBB: Blood-brain Barrier
EC: Endothelial Cells
EPL: Ependymal-like
FC: Fold-change
GSEA: Gene Set Enrichment Analysis
HUVEC: Human Umbilical Vein Endothelial Cells
IN: Inhibitory Neuron
IPC: Intermediate Progenitor Cell
MLC: Mesodermal-like Cell
OPC: Oligodendrocyte Precursor Cells
PC: Proliferating Cell
TRH: Thyrotropin-releasing Hormone
UMAP: Uniform Manifold Approximation and Projection
UPRC: Unfolded Protein Response Cells

## Supplementary figure legends

**S1 Fig. The analysis flow of multiple scRNA-seq data.**

(A) Analysis method overview. Cell types were individually assigned and integrated for intravascular organoids, non-vascular organoids, and fetal brains collected from public databases. (B) Annotation tree indicating the cell type assignment strategy. If there is no expression of the indicated marker gene, follow the path of “Negative” to be a candidate for the cell type below.

**S2 Fig. Pre-processing and integration of scRNA-seq data.**

(A) Parameters for each sample. Left panel: amount of characteristic RNA expression; middle panel: total RNA expression; right panel: mitochondrial expression. (B) Plot of standard deviation for each feature dimension to select the feature dimension with the smallest change in standard deviation. (C) Cell counts for each cell type in each sample. (D) Clusters of integrated samples. (E) UMAP for each integrated sample. (F) UMAP indicating cell types in each integrated sample. (G) Expression of each marker gene.

**S3 Fig. Characteristic gene expression profiles of annotated cell types.**

(A) Expression specificity of 100 genes characteristic of the cell type in each sample. Differentially expressed genes in each cell compared to other cell types were calculated, and the top 100 genes were sorted in descending order of log_2_FC with a cut-off at corrected p-value < 0.01. Each cell type has a characteristic gene expression pattern. (B) Overlap of characteristic genes for each cell type between each organoid and corresponding to the fetal brain. Differentially expressed genes specific to each cell group were identified with a cut-off at corrected p-value (). The genes overlapping with the Venn diagram were plotted using the “ggvenn” package (v0.1.9).

**S4 Fig. Evidence for cell type annotation and GO terms.**

(A–H) Expression of marker genes corresponding to cell types in each sample. (I) Plot of the differential expression levels of the gene sets characteristic of each GO term, visualized by the “heatplot” function of the “enrichplot” package (v1.16.2). The “ENTREZID” were converted to gene symbols using the “setReadable” function of the “DOSE” package (v3.22.1).

**S5 Fig. scRNA-seq data analysis in ectoderm cell subtypes.**

(A) UMAP in each sample of extracted neurons (IPC, EX, IN). (B) Alterations in GO-terms induced by vascularization in neurons. Note that color indicates enrichment score, not p-value. (C) Differentiation trajectories of ectodermal cells revealed by trajectory analysis.

**S6 Fig. Evidence for determination of MLC subtypes.**

(A) Eleven clusters of resolution = 0.3 obtained unbiased. The resolution was determined by the “clustree” shown in S6B Fig. For each cluster, a cell type was assigned based on the evidence in S6C and S6D Fig. (B) Clustree plot representing cluster relationships at resolutions from 0.1 to 1 (step 0.1) using the “clustree” package (v0.5.0). The dot size indicates the number of cells, and the line extending from the cluster indicates the cluster relationship. (C) Expression levels of genes that were characteristically expressed in each cluster. The color of the plots matches the concept color of each cell type. (D) GSEA results calculated based on non-biased computed gene sets. The table on the right side presents the p-value for each GO term. The plots were generated by the “gseaplot2” function of the “enrichplot” package.

## Supplementary tables

**S1 Table. Genes commonly upregulated by vascularization and their corrected p-values and average log_2_-fold changes are listed.**

The “upregulated” tab lists commonly upregulated genes and the “downregulated” tab lists commonly downregulated genes.

**S2 Table. List of genes altered by vascularization.**

No cut-off p-values are used in this table.

**S3 Table. List of genes altered by vascularization in neuronal populations.**

No cut-off p-values are used in this table.

**S4 Table. List of genes altered by vascularization in endothelial populations.**

No cut-off p-values are used in this table.

**S5 Table. Differentially expressed genes in the fetal brain of endothelial populations.**

No cut-off p-values are used in this table.

**S6 Table. Differentially expressed genes were specifically expressed for each cluster in the MLC population.**

No cut-off p-values are used in this table.

