## Supplementary figures and images for "Integrative analysis of *in vitro* strategies to induce human vascularized cerebral organoids using single-cell RNA sequencing data"

### S1 Fig

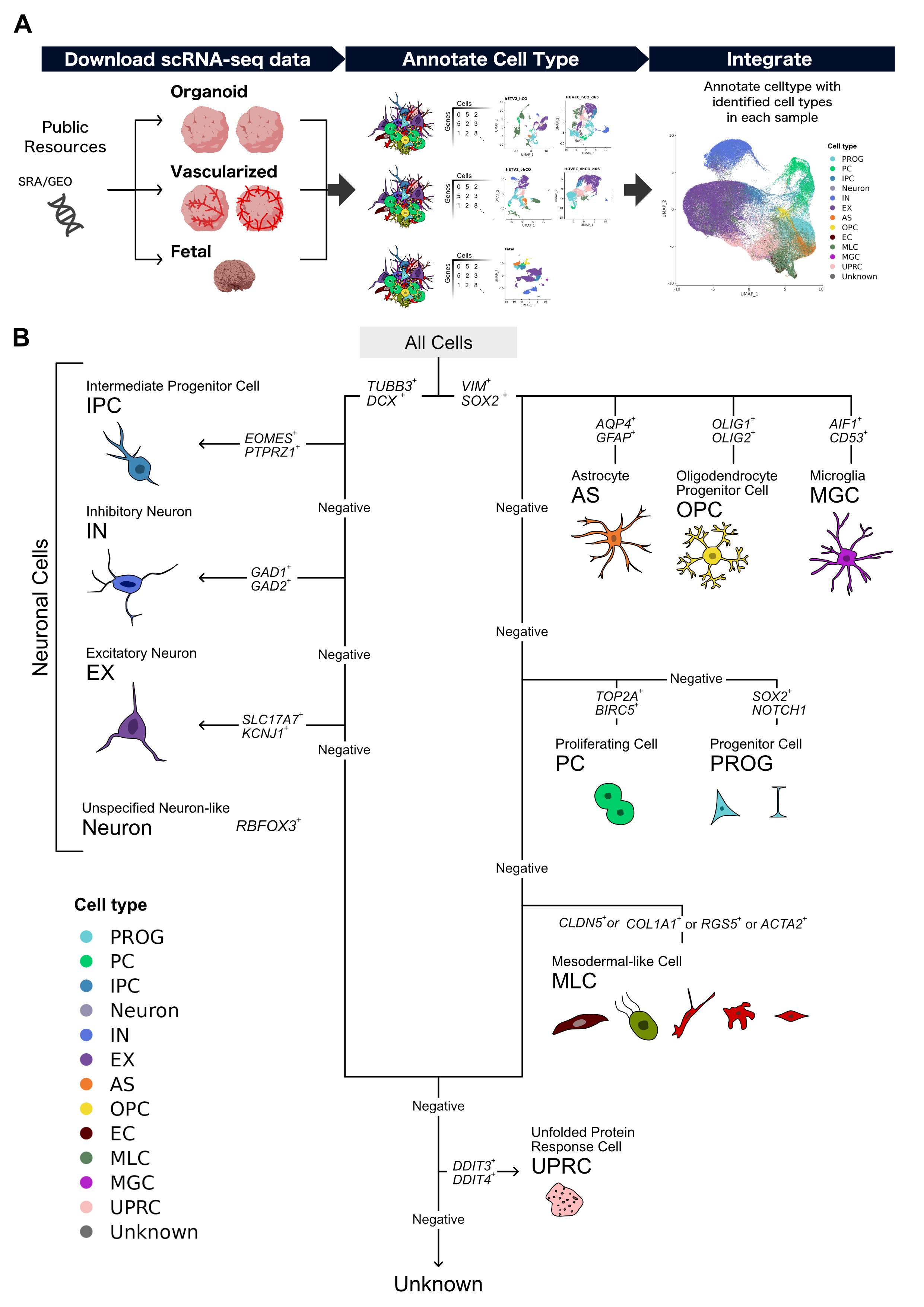

### S2 Fig

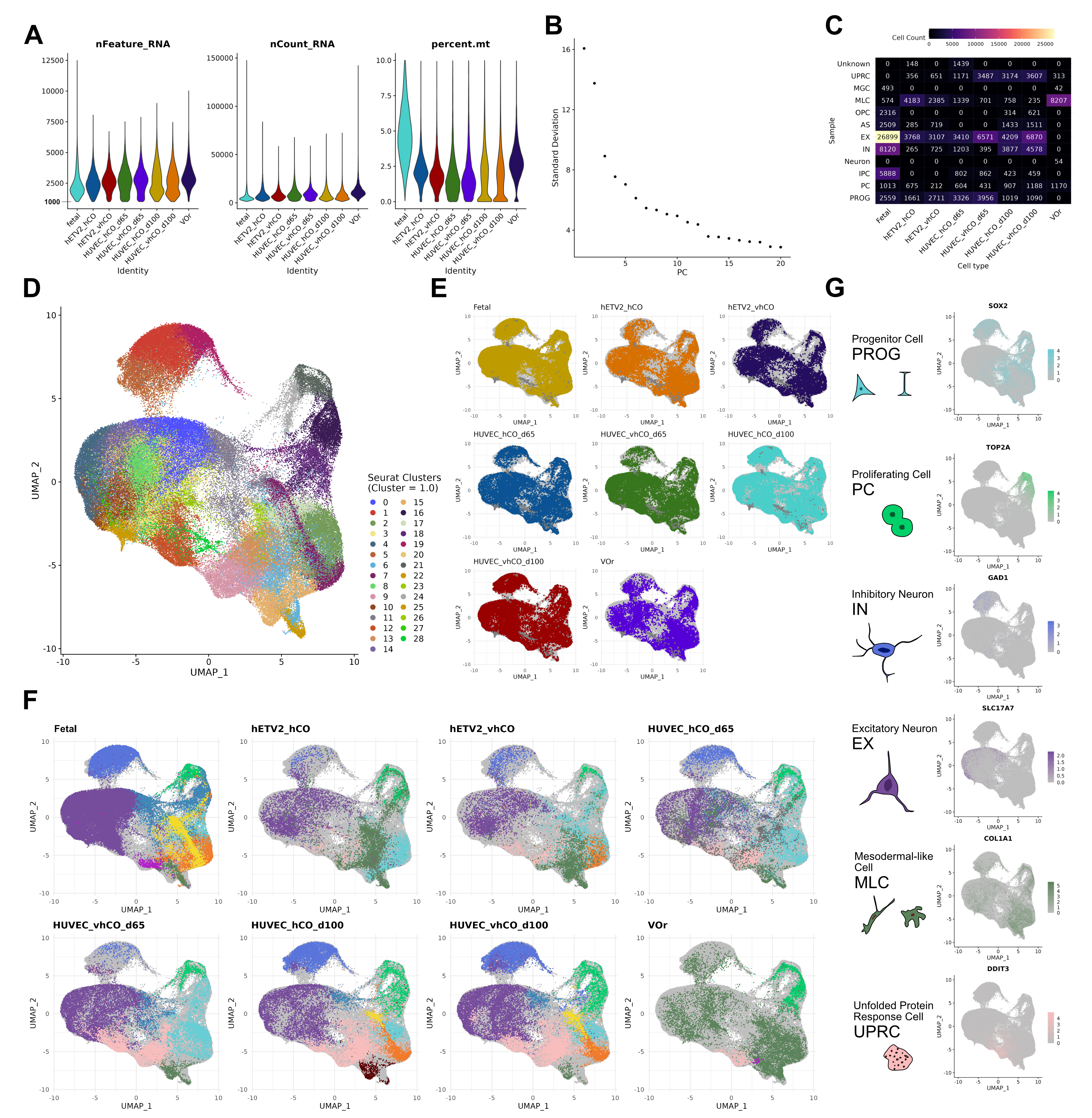

### S3 Fig

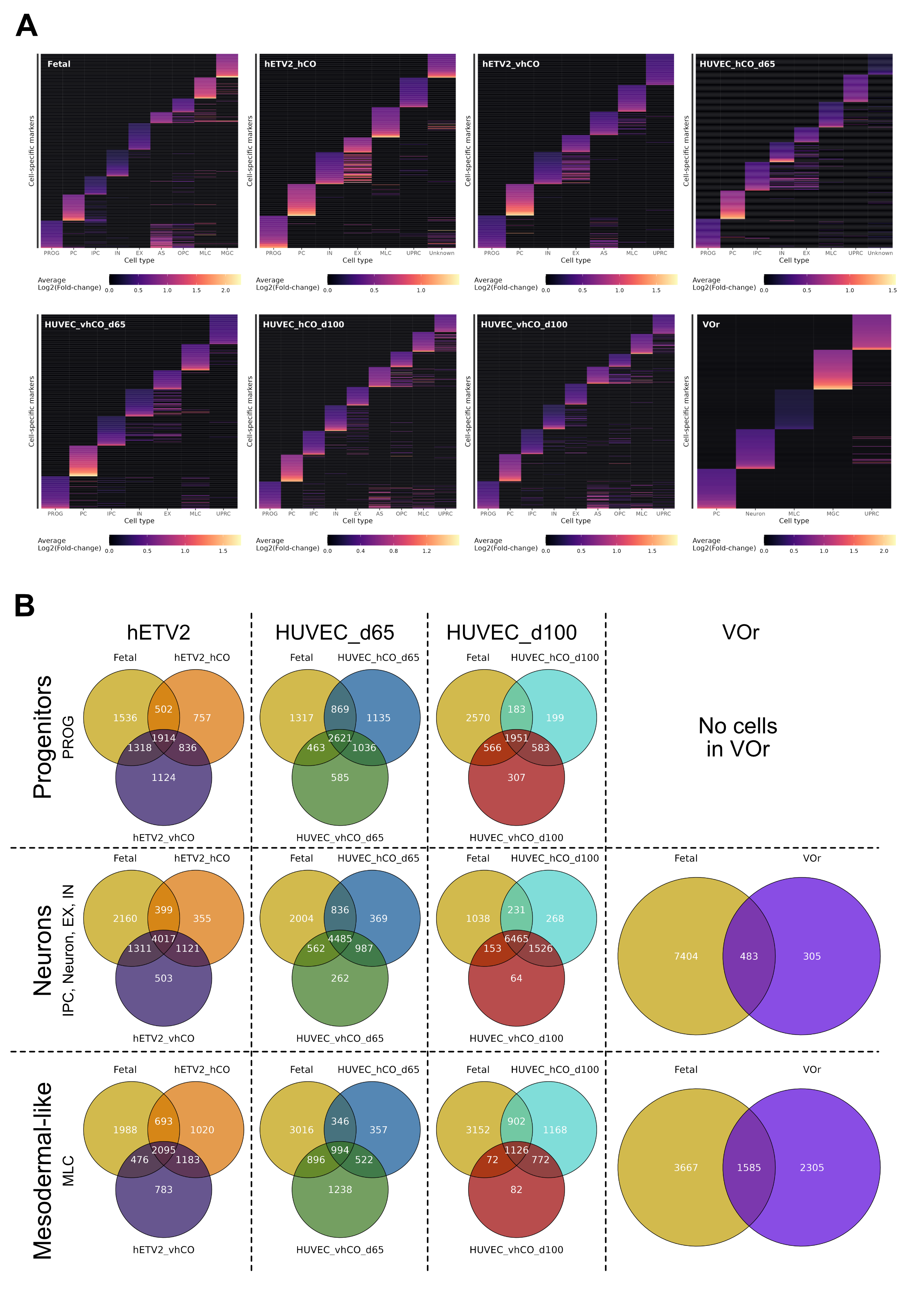

### S4 Fig

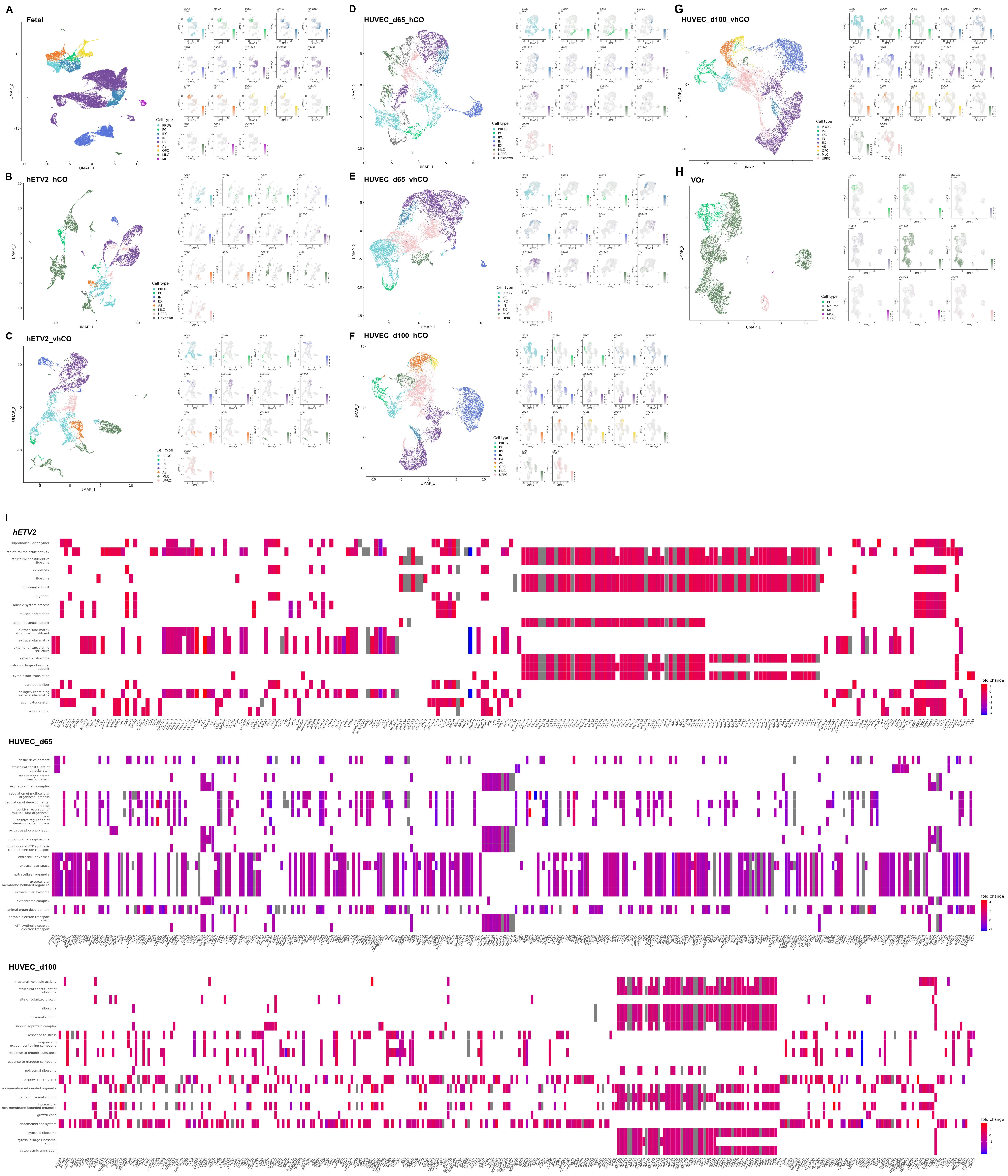

### S5 Fig

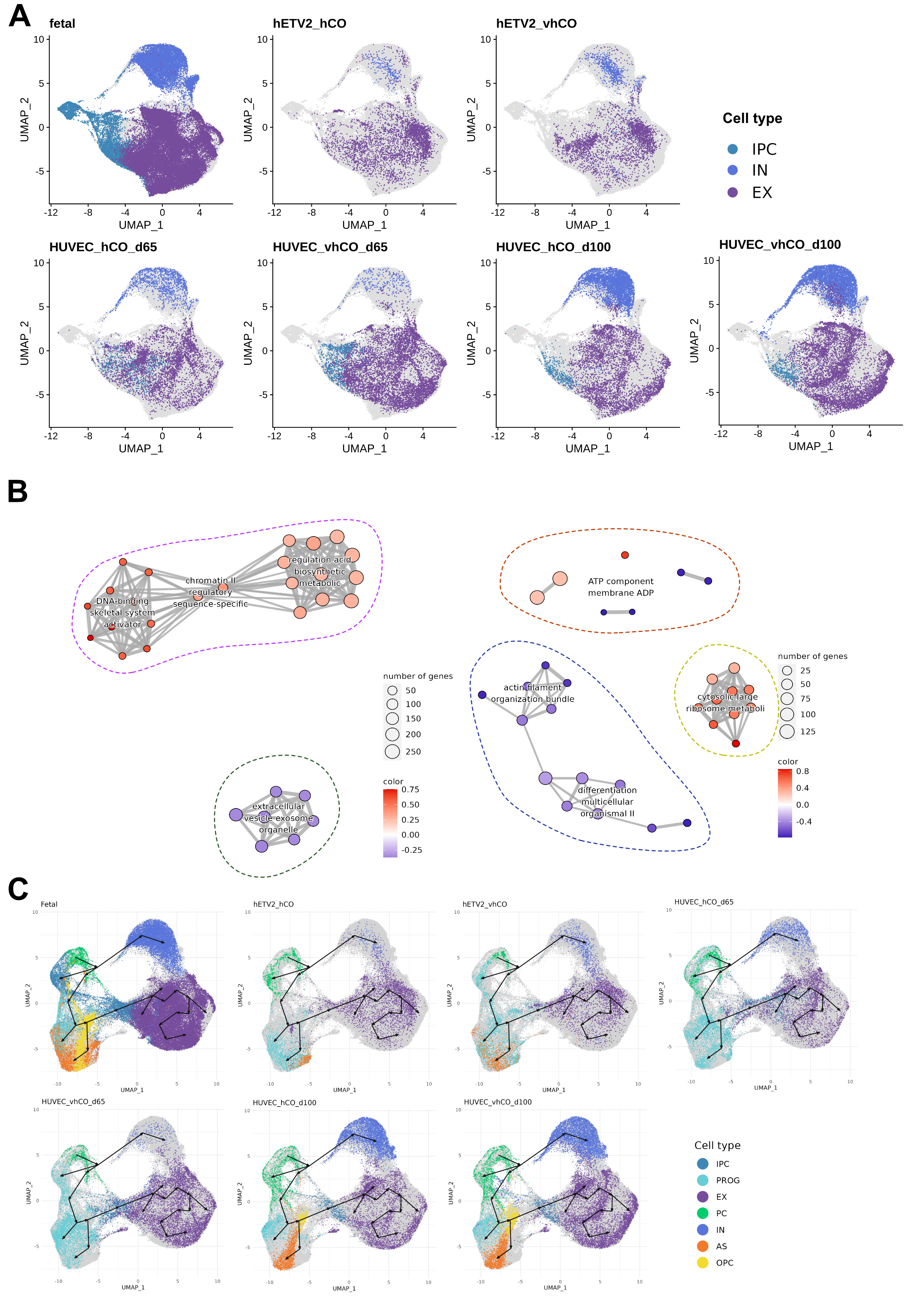

### S6 Fig

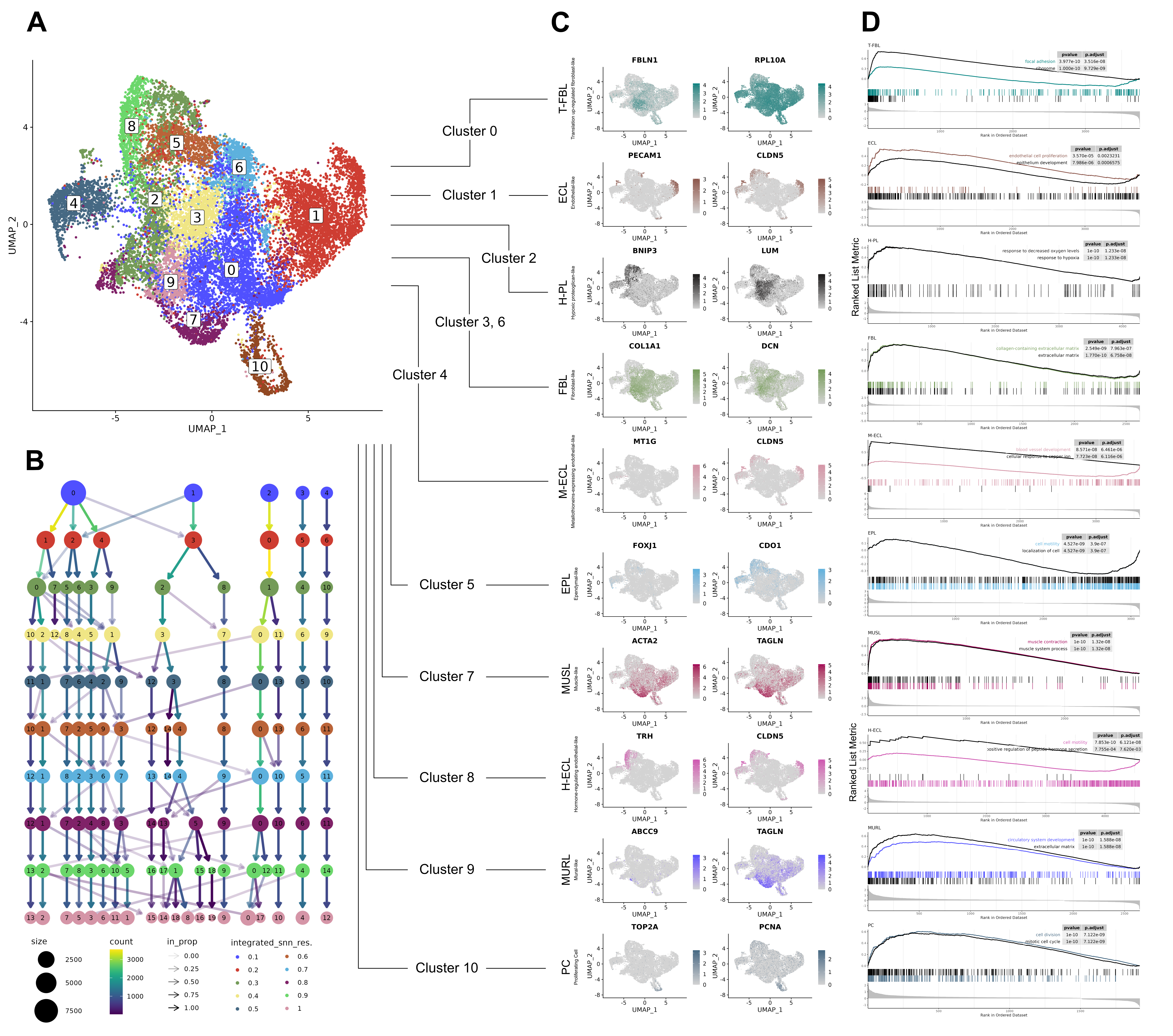
